# Using Y-chromosome capture enrichment to resolve haplogroup H2 shows new evidence for a two-Path Neolithic expansion to Western Europe

**DOI:** 10.1101/2021.02.19.431761

**Authors:** Adam B. Rohrlach, Luka Papac, Ainash Childebayeva, Maïté Rivollat, Vanessa Villalba-Mouco, Gunnar U. Neumann, Sandra Penske, Eirini Skourtanioti, Marieke van de Loosdrecht, Murat Akar, Kamen Boyadzhiev, Yavor Boyadzhiev, Marie-France Deguilloux, Miroslav Dobeš, Yilmaz S. Erdal, Michal Ernée, Marcella Frangipane, Miroslaw Furmanek, Susanne Friederich, Emmanuel Ghesquière, Agata Hałuszko, Svend Hansen, Mario Küßner, Marcello Mannino, Rana <u>Ö</u>zbal, Sabine Reinhold, Stéphane Rottier, Domingo Carlos Salazar-García, Jorge Soler Diaz, Philipp W. Stockhammer, Consuelo Roca de Togores Muñoz, K Aslihan Yener, Cosimo Posth, Johannes Krause, Alexander Herbig, Wolfgang Haak

## Abstract

Uniparentally-inherited markers on mitochondrial DNA (mtDNA) and the non-recombining regions of the Y chromosome (NRY), have been used for the past 30 years to investigate the history of humans from a maternal and paternal perspective.

Researchers have preferred mtDNA due to its abundance in the cells, and comparatively high substitution rate. Conversely, the NRY is less susceptible to back mutations and saturation, and is potentially more informative than mtDNA owing to its longer sequence length. However, due to comparatively poor NRY coverage via shotgun sequencing, and the relatively low and biased representation of Y-chromosome variants on capture arrays such as the 1240K, ancient DNA studies often fail to utilize the unique perspective that the NRY can yield.

Here we introduce a new DNA enrichment assay, coined YMCA (Y-mappable capture assay), that targets the “mappable” regions of the NRY. We show that compared to low-coverage shotgun sequencing and 1240K capture, YMCA significantly improves the coverage and number of sites hit on the NRY, increasing the number of Y-haplogroup informative SNPs, and allowing for the identification of previously undiscovered variants.

To illustrate the power of YMCA, we show that the analysis of ancient Y-chromosome lineages can help to resolve Y-chromosomal haplogroups. As a case study, we focus on H2, a haplogroup associated with a critical event in European human history: the Neolithic transition. By disentangling the evolutionary history of this haplogroup, we further elucidate the two separate paths by which early farmers expanded from Anatolia and the Near East to western Europe.

## Introduction

Uniparentally inherited markers such as mtDNA and the NRY are an attractive source of information about the demographic history of a population due to the fact that their history can be represented by a simple evolutionary tree (Brown 1980; Jobling et al. 2003). Since the seminal studies of the 1980s (Cann et al. 1987; Torroni et al. 1984) and prior to the genomic era, much of the genetic history of humankind and the peopling of the world was inferred from uniparentally inherited mtDNA and NRY (e.g. Pakendorf and Stoneking 2005; Underhill et al. 2007; Kivisild 2017)

Due to the high copy number of mtDNA in the cells (Ingman and Gyllensten 2001), the short genome length (<17kB), and the relatively high substitution rate (Soares et al. 2009), mtDNA has been particularly well-studied, yielding an inexpensive and yet reliable source of information about the genetic variability of a population (Finnilä et al. 2001; Torroni et al. 2006; Posth et al. 2016).

Conversely, the mappable portion (the regions for which short reads, such as in ancient DNA studies, have been reliably mapped) of the NRY is much longer (~10,445kB) and presents only as single-copy in the cells of male individuals. The estimated evolutionary substitution rate is up to two orders of magnitude lower for the NRY (2.5 × 10^-9^) (Helgason et al. 2015) than for the coding regions of mtDNA (~2.2 × 10^-7^) (Soares et al. 2009; Fu et al. 2014; Posth et al. 2016). However, the greater genome length of the NRY, compared to the mtDNA, means that from these numbers, and for a single lineage, we still expect to observe a point mutation approximately every 109 to 130 years for the NRY, compared to between 262 to 7360 years for mtDNA. Consequently, the NRY can contain more information about the paternal demographic history of a population and can be informative about male-biased population demographic changes, such as through male-driven migration (Karmin et al. 2015; Zeng at al. 2018; Olalde et al. 2019), or patrilocality (Deguilloux et al. 2013) so seeking insights into the paternal history of a population can be of critical importance.

When studying the demographic history of humans, aDNA has been shown to be an irreplaceable source of information. aDNA studies have revealed large-scale population movements and genetic turnover events in Western Eurasia (e.g. Fu et al. 2016; Allentoft et al. 2015; Haak et al. 2015; Olalde et al. 2018) that were otherwise impossible to recover from modern data. Studies of the uniparentally inherited markers of ancient individuals have also yielded otherwise undetectable results, e.g. the loss of European mtDNA diversity following the repeopling of Europe after the last glacial maximum (Posth et al. 2016), or the decrease in and partial replacement of diversity of hunter-gatherer Y-chromosome lineages in eastern and central Europe following the Neolithic expansion (Bramanti et al. 2009; Haak et al 2010; Brandt et al. 2013; Lipson et al. 2017; Mathieson et al. 2018), followed by the loss of diversity of Neolithic Y-chromosomes lineages with the arrival of Steppe-like ancestry at the beginning of the 3^rd^ millennium BCE (Allentoft et al. 2015; Haak et al. 2015; Olalde et al. 2018, 2019).

Researchers using aDNA data usually encounter problems related to sample quality, specifically a decrease of endogenous human DNA due to post-mortem DNA decay and environmental contamination (Higgins et al. 2015). The Y chromosome makes up <2% of the total DNA in male cells, meaning that if researchers wish to use shotgun (SG) sequencing to adequately cover enough informative sites on the single-copy NRY, then, even for samples with good DNA preservation, a substantial sequencing effort is required.

The development of targeted capture arrays has allowed aDNA researchers to enrich specific sites and regions of the genome for sequencing, vastly improving the yield of human endogenous DNA from ancient samples (Fu et al. 2014; Mathieson et al. 2015). One such popular array is the 1240K array, which targets ~1.24M ancestry-informative sites on the human genome, of which ~32K represent a selection of known variants on the Y chromosome (based on an ISOGG list of informative Y-chromosomal SNPs as of 2013/14)(Mathieson et al. 2015).

This relatively low number of targeted Y-SNPs, compared to the number of currently known, informative Y-SNPs (as defined by ISOGG, n=73,163, or Yfull, n=173,801, https://isogg.org/tree; https://www.yfull.com), allows for basic Y haplogroup (YHG) assignments, but is heavily biased towards modern-day diversity and certain geographic regions. As a consequence, depending on the representation of particular Y-SNPs on the 1240K array, the resulting YHG assignments can be of low and uneven resolution.

To better study and understand the male history of human populations, we saw a need for a targeted array that specifically enriches sequence data for sites on the NRY, without targeting only already well-known SNPs. To achieve this, we designed and implemented YMCA (Y-mappable capture assay), a tiled capture-array for NRY sequence data that targets regions of the NRY for which short reads, typical in ancient DNA samples, are reliably mapped to the human genome, as defined by Poznik et al. (2013). A similar approach has been explored by two previous studies (Cruz-Dávalos et al. 2018; Petr et al. 2020). However, we avoid in-depth comparison with the probe set presented by Petr et al. which targets ~6.9 MB was designed to substantially older samples, such as Denisovan and Neanderthal individuals, and hence the definition of “mappable” was far more conservative and stricter. Conversely, Cruz-Dávalos et al. also present a capture-enrichment approach designed for ancient human samples with low endogenous DNA. The reported ~8.9 MB regions are almost completely included in our target regions (99.97%), and we show that the remaining ~1.5 MB in our target regions still yield reliably mapped sites (see Supplementary Table S1.4).

Here we show that YMCA significantly improves the relative coverage of NRY sites when compared to shotgun sequencing, allowing for the enrichment of NRY sites for the same sequencing effort. We also show that YMCA significantly outperforms 1240K SNP array sequencing in two ways. Empirically, we show that YMCA improves the number of NRY sites that are hit. We also show that, by considering the targeted NRY sites as defined by the associated bed files, that if we were to sequence a sample with high complexity to exhaustion, that YMCA has an improved potential resolution for Y-haplogroup assignment and the discovery of new diagnostic SNPs when compared to 1240K array sequencing.

We highlight the improved performance obtained via YMCA by analysing the Y-chromosomal haplogroup H2 (H-P96), a low-frequency YHG that is associated with early farmers during the Neolithic transition in Western Eurasia. We curated a data set of 46 previously published individuals (45 ancient and 1 modern), and 49 newly YMCA-sequenced individuals (all ancient). We show that our current understanding of H2, which is based largely on modern H2 samples (n=20), is inconsistent with the ancient diversity of our H2 individuals. In resolving this ancient haplogroup, we can show two distinct migration paths along the Mediterranean and Danube for Neolithic groups from Anatolia to Western Europe, ultimately resulting in the Mediterranean-derived groups also reaching the Atlantic Archipelago/Britain and Ireland/British Isles.

## Results and Discussion

### Validating the performance of YMCA

To evaluate the performance of our new NRY-Capture assay (YMCA), we calculated the empirical fold-increase in endogenous human DNA for a range of samples with varying levels of preservation. We then compared the empirical performance of YMCA against standard shotgun sequencing and 1240K capture on the same libraries by inspecting the number of NRY sites hit, as well as the number of ISOGG SNPs hit at least once for each library type. We account for sample quality and input sequencing effort by filtering for only human endogenous reads, and then normalising the number of sites/SNPs hit per five million endogenous reads.

We observed a significant fold-increase in the amount of endogenous human DNA when comparing shotgun sequencing to YMCA (see Figure S4), which we refer to as “enrichment” from here on. We found that enrichment diminished as the preservation of the sample increased, i.e. for samples with higher starting endogenous DNA % the effect of the enrichment was reduced, but still significant.

We observed a significant mean fold increase of ~15.2X in the number of NRY sites hit with YMCA captured libraries when compared to shotgun sequencing (p=5.5 × 10^-7^), and ~1.84X when compared to 1240K sequencing (p=8.8 × 10^-12^), showing that YMCA hits on average more NRY sites than both shotgun and 1240K sequencing. This also indicated that, since we hit on average 15.2 times as many ISOGG SNPs per five million reads for SG sequencing, we would need to sequence ~76 million reads to hit the same number of NRY sites for shotgun sequencing compared to only 5 million reads for YMCA.

Interestingly, we also found mean fold increase of ~4.36X in the number of ISOGG SNPs hit with YMCA captured libraries when compared to 1240K sequencing (p=9.0 × 10^-14^), indicating that, for the same sequencing effort, YMCA also hits more informative SNPs, for the same sequencing effort.

We also found that the fold-increase in the number of NRY sites that we hit, and the endogenous DNA percentage for shotgun and 1240K sequencing were uncorrelated (p=0.976 and p=0.617), and that the number of ISOGG SNPs hit and the endogenous DNA percentage for 1240K sequencing were uncorrelated (p=0.1825) indicating that our results are not dependent on sample quality. Hence, we found that, although the SNPs hit on the Y chromosome are an added bonus when using the 1240K array, as it is primarily used for analysing the autosomal genome of male and female individuals, YMCA is clearly a significant improvement if researchers wish to efficiently and thoroughly investigate the non-recombining portion of the Y chromosome.

We then compared the percentage of haplogroup-informative SNPs on the ISOGG SNP list v14.8, that are also included in the 1240K array, and YMCA, according to their respective bed files. This comparison will be particularly powerful as YMCA and the 1240K array are based on the same technology, and captured via identical lab protocols. The 1240K array targets 24.44% of the currently listed ISOGG SNPs, whereas the YMCA targets 90.01% (Figure 1). Note that the remaining 9.99% of ISSOG SNPs exist in regions of the NRY which are considered “unmappable” for short reads common in ancient DNA. Since each of the sites in the1240K array is targeted by two probes (allele and alternate allele) and two 52bp probes on either side of the variant, additional sites flanking the “targeted” sites can also be recovered from the mapped reads. Hence, we also allow a window of 120bp (60bp on either side) for each SNP on the 1240K array, which is a reasonable average read length for aDNA. For this 1240K+60bp list of sites the percentage of targeted ISOGG SNPs increases to 45.34%, but this also illustrates that the 1240K SNP array is fundamentally limited by the total number of informative Y chromosome SNPs included.

**Figure 1:**
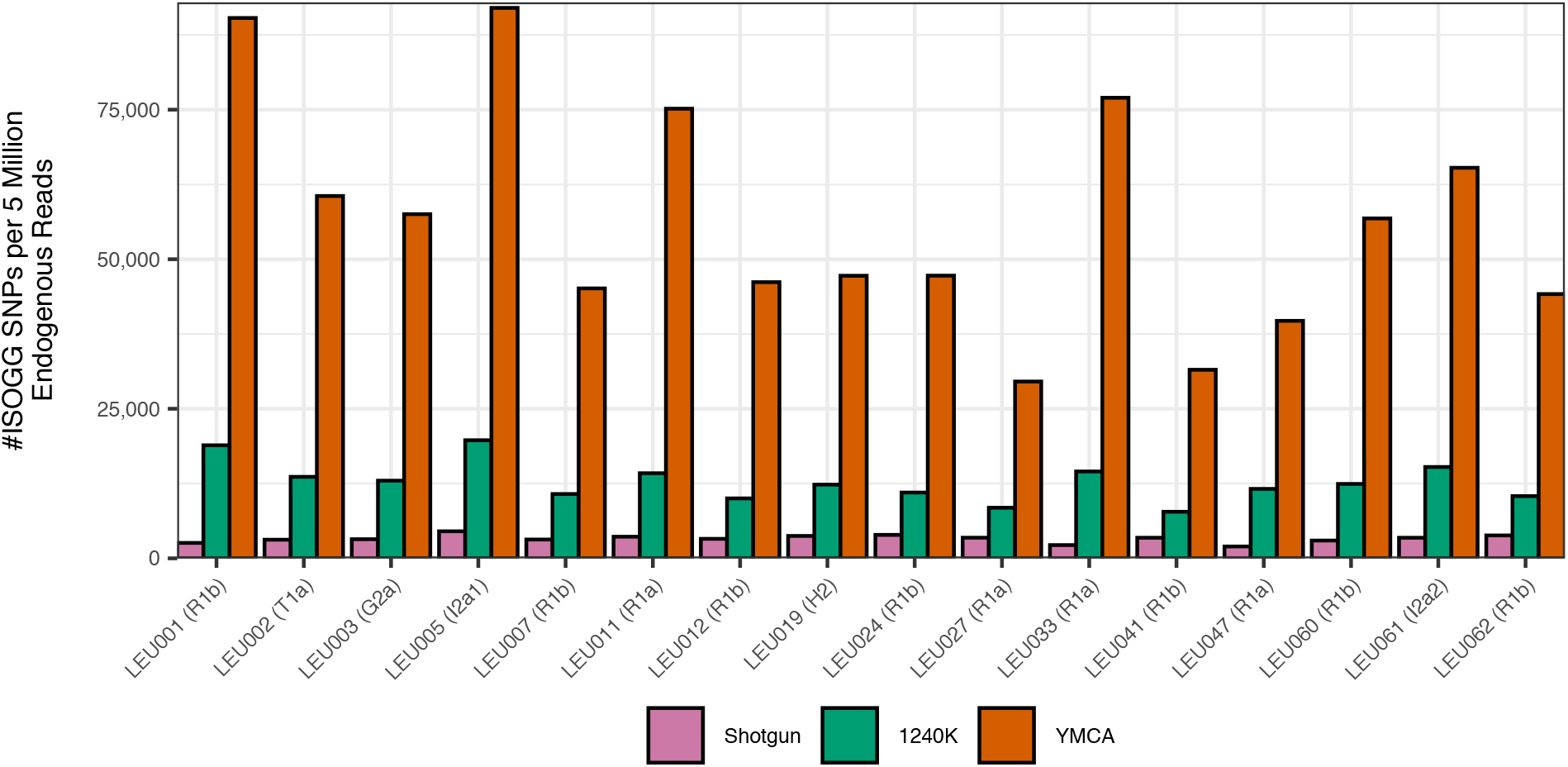
The number of mappable non-recombing Y chromosome (NRY) sites hit per 5M quality-filtered mapped reads (y-axis) for the same libraries (x-axis) for shotgun, 1240K and YMCA sequencing (colours).

Additionally, recovering as much of the NRY as possible is of critical importance, especially when researchers are interested in looking for new variants on the Y chromosome, or uncovering past diversity that might no longer exist. When comparing the raw number of sites targeted by the 1240K array to YMCA, we observed that the 1240K capture array potentially targets a total of 32,670 sites, which is approximately 0.31% of the number of SNPs targeted by YMCA (~10,445kB). However, if one is to include a window of 120bp around each SNP again, then the 1240K array potentially targets ~3,953kB sites or 37.82% of the number of sites one can potentially analyse using our YMCA. Hence, YMCA is a predictably better tool for exploring the NRY for new ancestry informative SNPs.

We were also interested in comparing the potential resolution to which YHG assignments can be made, given the available ISOGGs SNPs targeted by YMCA and 1240K. We also found that the resolution of a YHG call cannot be improved, even when including a 120bp tiling window around the ~32K Y-SNPs of the 1240K array, according to the ISOGG SNPs occurring in the respective bed files. This holds true both for dominant YHGs today and in particular for those that are associated with known ancient populations, but that have significantly reduced in frequency in modern populations, and which are not well covered for diagnostic SNPs on the 1240K array.

We often observe low resolution in haplogroups such as the early hunter-gatherer haplogroup CV20 (Fu et al. 2016), and the Neolithic expansion-associated G-Z38202 (Lacan et al. 2011) and H-P96, for which the Y-SNPs of the 1240K array target 0.8%, 0% and 13% of the associated ISOGG SNPs, respectively (we include SNPs within three branches downstream of each terminal SNP). If we include a 120bp window, then these percentages increase to a more respectable 32.5%, 31.2% and 36.2%, which are still much lower than the 89.6%, 90.6% and 95.2% of SNPs targeted by YMCA (see Figure 2). In addition, poor theoretical coverage for YHGs which are thought to be present in early human population movements, but which remain relatively prevalent in modern populations, can still be an issue for sites on the 1240K array. For example, Q-M3, which is associated with the initial peopling of the Americas (Ruiz-Linares et al. 1999) has only 11.9% (33.5% if a 60bp window is included) of the relevant diagnostic SNPs covered, compared to 92% for YMCA.

**Figure 2:**
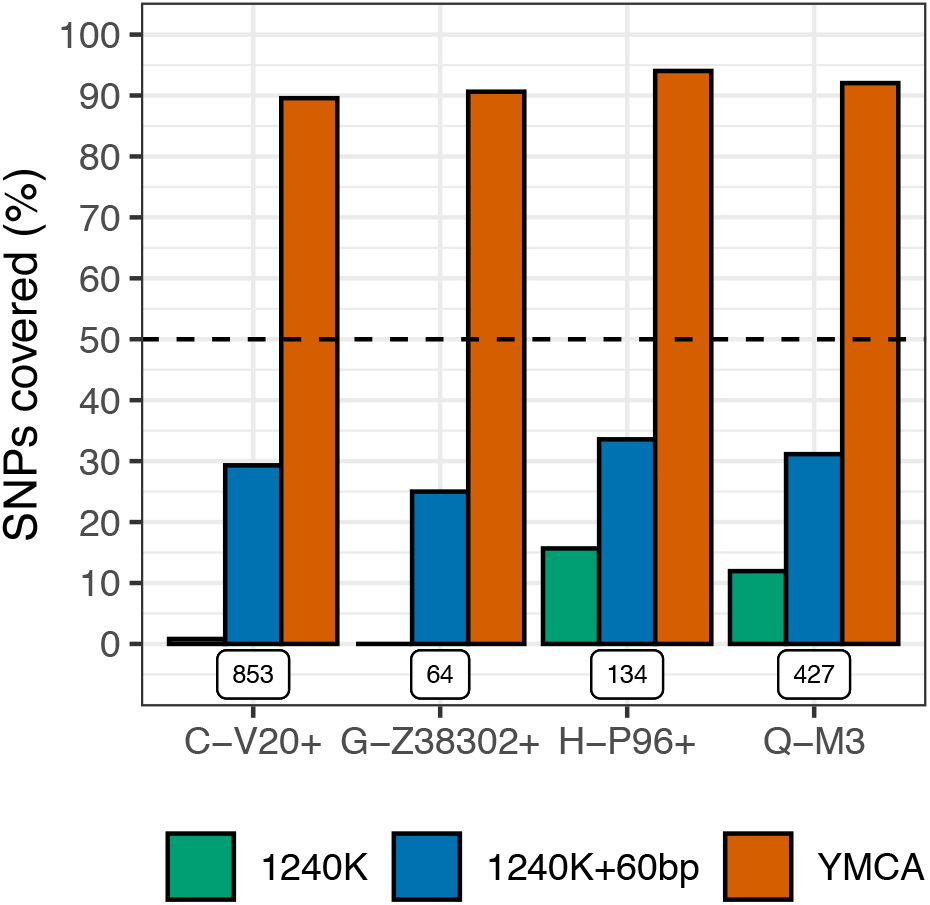
The percentage of SNPs (y-axis) covered (up to three branches downstream) for four Y-chromosome haplogroups (x-axis) associated with ancient populations. Colours indicate array SNPs targeted for 1240K (green), 1240K with a 60bp window (blue) and our YMCA (orange). The dashed black line indicates at least half of the SNPs are represented, and the total number of targetable SNPs is given below each group.

To summarize, YMCA enriches the relative proportion of reads mapping to the NRY, when compared to shotgun sequencing and the 1240K array. YMCA also targets more than 2.5 times as many sites on the NRY than the 1240K array, allowing for the detection of new diagnostic SNPs. Critically however, YMCA targets SNPs which are already known to be informative, but the 1240K array cannot target.

### Application of YMCA to YHG H2 as a case study

Through routine application of SG sequencing for sample screening, followed by 1240K capture for suitable samples in our lab, we were able to explore the performance of the new YMCA on a range of YHG in ancient male individuals. Here, we showcase an example of YHG H-P96, for which the resolution of the evolutionary tree is still unclear due to the scarcity of data and low frequency in modern-day populations. Judging from our current ancient DNA record, it appears that YHG H was more common in the past, in particular among males that were associated with the spread of farming across West Eurasia during the Neolithisation. As a result, we can show that aDNA research, and in particular high-resolution typing of YHG, can help elucidate the evolutionary relationship of Y chromosome lineages past and present.

The YHG H (H-L901) is thought to have formed in South Asia approximately ~48 kya (Sengupta et al. 2006). Three subsequent sub-haplogroups, H1 (H-M69), H2 (H-P96) and H3 (H-Z5857), appear to have quickly formed over the following four thousand years. H1 and H3 have estimated formation times of ~44.3 kya, however, H2 is estimated to have formed slightly earlier at ~45.6 kya [https://www.yfull.com].

H1 and H3 are still found in frequencies as high as 20% in South Asia (Rai et. al. 2012), but in extremely low frequencies in Europe, with H1 only being found associated with the spread of the Romani people ~900 ya. Conversely, H2 has been present in Europe since at least 10 kya (Lazaridis et al. 2016), and is strongly linked with the spread of agriculture (Hofmanová et al. 2016; Rivollat et al. 2020), but is found at no higher than 0.2% frequency in modern-day western European populations. In contrast, H2 was more common in Neolithic groups, and has been found to have constituted between 1.5% and 9% of the observed Y haplogroups, with the exception of the highly related samples from Rivollat et al., for which H2 was ~30% (Brunel et al. 2020, Haak et al. 2015, Mathieson et al. 2015, Lazaridis et al. 2016, Lipson et al. 2017, Olalde et al. 2019, Rivollat et al. 2020, Skourtanioti et al. 2020).

With the arrival of Steppe-related ancestry ~5 kya, incoming YHGs such as R1a and R1b would largely replace many of the older, Neolithic YHGs, such as G2, T1a, and H2 (Haak et al. 2015), and although H2 was never found in particularly high frequencies among Neolithic individuals, we expect that its diversity was also greatly reduced, and many sub-lineages were potentially lost altogether.

To test whether our YMCA could improve the haplotyping quality to a point which would allow us to also draw phylogeographic inferences, we made use of newly collated collection of prehistoric ancient human DNA data and selected individuals, who were tentatively assigned to YHG H2. While H2 is commonly found alongside the more dominant Neolithic YHG G2a (G-Z38302) (Lipson et al. 2017, Hofmanova et al. 2016), it is precisely the low frequency of H2 which is of interest here. The relative scarcity of H2 individuals, especially compared to the relatively high frequency of the accompanying G2a individuals, allows us to better track the ‘genealogical history’ and thus potential dispersal routes as we would expect a stronger effect of lineage sorting and therefore a higher chance of observing geographic patterns. In this particular case, we could trace expanding Neolithic farmers from Anatolia to Western Europe through the use of unique markers associated with H2 individuals and test whether we can genetically discern the proposed so-called “Danubian or inland’’ and “Mediterranean’’ routes of the Neolithic expansion (Price 2000), which had recently also found support by genomic signals from the nuclear genome (Rivollat et al. 2020). Unfortunately, we found that the H2 subsection of the evolutionary tree for the Y chromosome is currently poorly understood (due to the scarcity of modern samples of H2 individuals and the relative rarity of ancient H2 individuals), and, in many cases, inconsistent with a tree-like history for almost all of the published and unpublished ancient samples. In all but one case that we found that H2 individuals carry a mixture of derived SNPs from two bifurcated clades in the current ISSOG topology, such as from H2a1 and H2b1. Encouraged by the performance of the YMCA presented above, we thus analysed further H2 individuals in an attempt to resolve the branching pattern of this lineage.

For a non-recombining portion of DNA, the evolutionary history is expected to follow a tree-like structure, and therefore hybrids of sibling haplogroups (such as between ISOGG H2a, H2b1 and H2c1a) are impossible. To try and find a better resolved evolutionary history for these individuals, we constructed a maximum likelihood (ML) phylogenetic tree using IQ-TREE (see Methods). We identified two major clades from the ML tree (see Figure 3B), and denoted these two tentatively named clades H2d (green clade) and H2m (blue clade). Within each of our clades, we could also identify two sub-clades, denoted H2d1, H2d2, H2m1 and H2m2. Low bootstrap support values or sub-clades based on sites within these sub-clades indicated that no further haplogroup resolution was possible with our data. With respect to the current ISOGG nomenclature, we note that H2m appears to be defined by a mix of H2, H2a, H2a1~ and H2c1a~ SNPs, with H2m1 defined by additional H2a and H2c1a~ SNPs. Conversely, H2m2 is defined by SNPs which were previously undetected, indicating that H2m2 may be an extinct lineage of H2. Similarly, H2d appears to be defined by a mix of H2, H2a1~ and H2b1 SNPs, but with no H2a SNPs, while sublineages H2d1 and H2d2 are defined by SNPs which were previously undetected.

**Figure 3:**
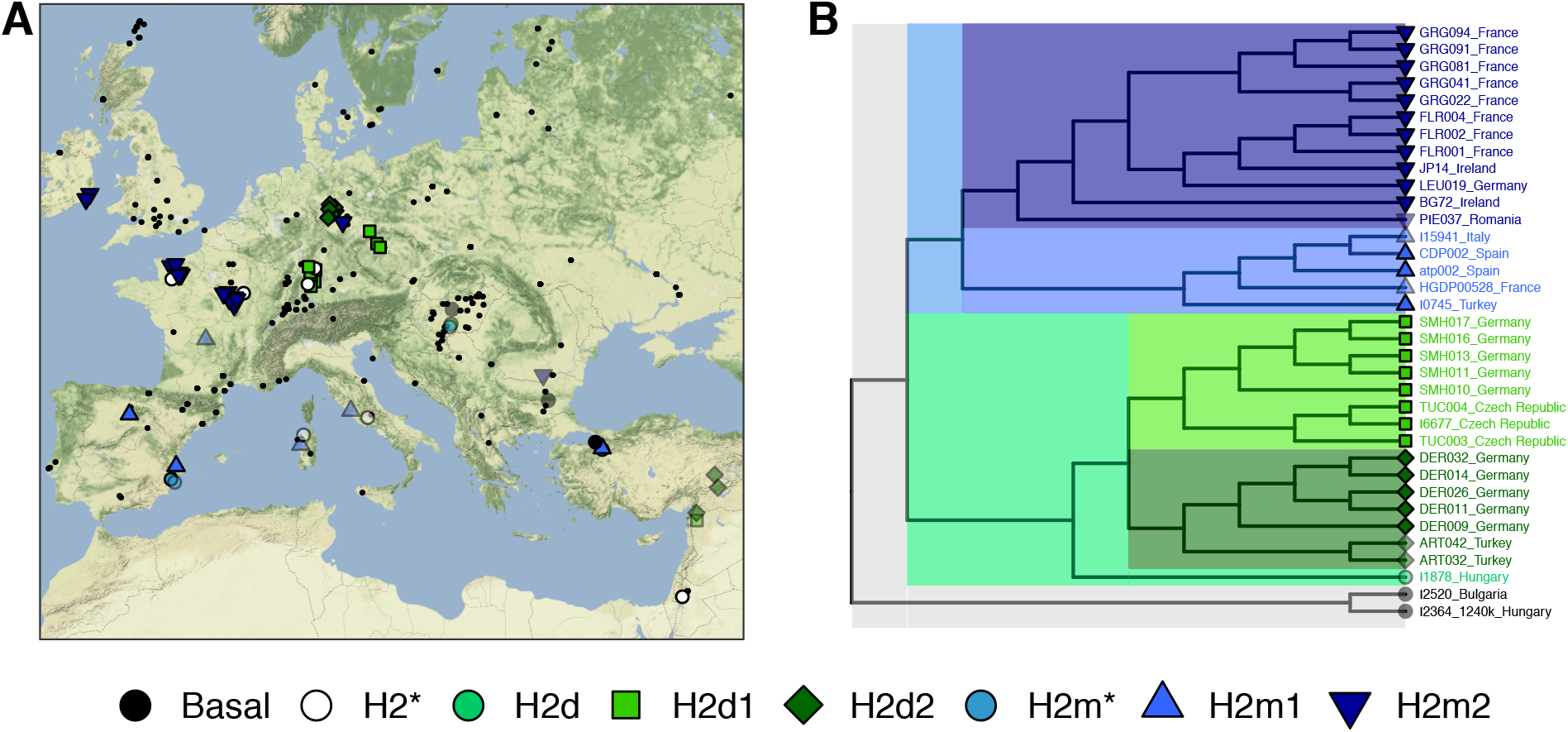
(A) H2 sample locations and (B) phylogenetic relationships (no branch length units). Shapes and colours indicate the four major clades inferred from the phylogenetic tree. Symbol shading indicates early to late Neolithic (solid) or post-Neolithic (transparent). Black dots indicate all non-H2 Neolithic individuals from Freeman2020 to indicate H2 sampling prevalence. Stars in haplogroup assignments in (A) indicate a lack of resolution to assign samples (not used in the ML tree) to downstream sub-haplogroups.

Based on our extended set of diagnostic SNPs, we were able to assign n=58 of our samples to either one of these four sub-clades, one of the two major clades, or (basal) H2* (due to low coverage) even for samples that did not meet minimum coverage requirements to be included in the ML tree. Interestingly, we found one individual, I1878, who was derived for our H2d SNPs, but ancestral for SNPs in H2d1 or H2d2. Finally, we also had three individuals who were not derived for any of these additional SNPs, and were ancestral for many of the H2 SNPs (denoted basal, n=3).

When we plotted all of the samples in our study on a map of Europe, a phylogeographic pattern clearly emerged. The H2d individuals are all found along the so-called inland/Danubian route into central Europe, and all but one of the H2m individuals are found along the so-called Mediterranean route into Western Europe, the Iberian Peninsula and ultimately, Ireland. The solitary H2m individual (LEU019) found in central Germany is dated to the Late Neolithic/Early Bronze Age context, postdating the Neolithic expansion by 2000-3000 years. Archaeological and mtDNA evidence of an eastward expansion of Middle/Late Neolithic groups such as Michelsberg (Jeunesse 1998; Küβner 2016, Beau et al. 2017) could potentially explain this single geographically outlying observation.

Due to the incomplete and varying coverage of our ancient samples, we were unable to reliably produce a calibrated tree for divergence time estimates using the radiocarbon dates of ancient samples as tip dates. Instead we estimated the time since the most recent common ancestor (TMRCA) for each pair of individuals to investigate the split times for our newly identified H2 clades (see Methods). First, we calibrated our relative substitution rate so that we estimated a mean TMRCA of ~161.3 kya for haplogroup A0 with all other haplogroups (see Figure S2). Using this calibrated substitution rate, we estimated a TMRCA for haplogroup A1 of ~133.2 kya, and a TMRCA of ~48 kya for haplogroup HIJK, which are extremely close to the current estimates of ~133.4 kya and ~48.5 kya respectively [https://www.yfull.com]. Our estimated TMRCA for H2 was ~24.1 kya, which is slightly older than the current estimate of ~17.1 kya, which could be explained by our extremely limited access (only one) to high-coverage modern H2 samples, and our increased number of ancient samples [https://www.yfull.com].

We found that the estimated TMRCA for H2d and H2m was ~15.4 kya. We also found that H2m1 and H2m2 had an estimated TMRCA of ~13.3 kya, and that H2d1 and H2d2 had an estimated TMRCA of ~11.9 kya (see Figure 4). We note, however, that even though the associated error bars are wider due to fewer overlapping SNPs, the mean estimates are still relatively consistent. These estimates, plus the fact that H2d1 and H2d2 individuals are found in Anatolia and the Levant, show that H2 diversity most likely existed in Near-Easterner hunter-gatherers before the establishment of agriculture and animal husbandry and likely also in early farmers, and subsequently spread via the Neolithic expansion into Central and Western Europe.

**Figure 4:**
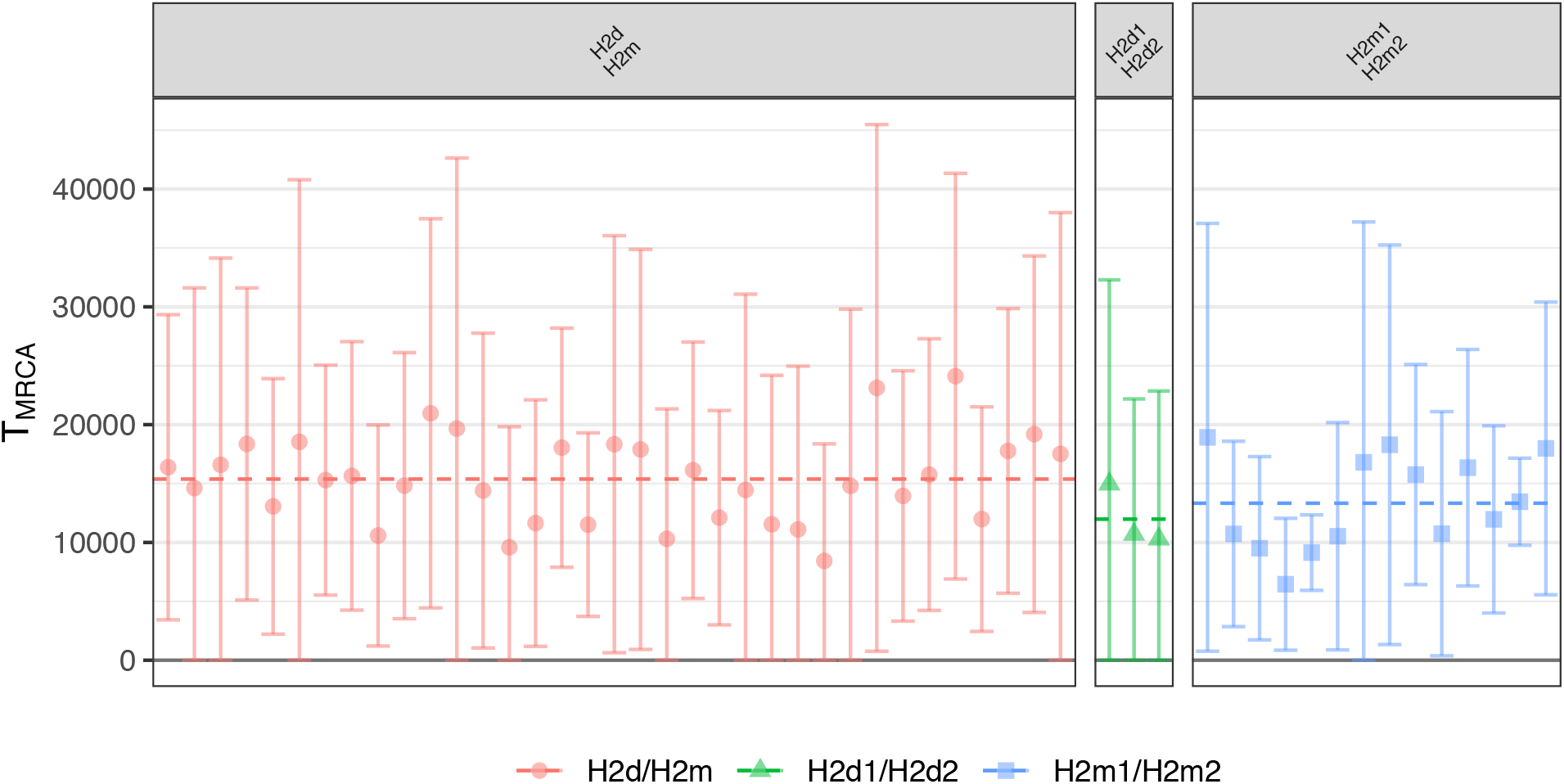
Estimated time since most recent common ancestor (T_MRCA_) (y-axis) for each H2 subgroup with each other (facets), calibrated by the split time of ~163 kya of haplogroup A0 with all other Y haplogroups. Error bars indicate 95% confidence intervals.

### Identifying diagnostic SNPs for improved YHG H2 resolution

Having used YMCA to identify four novel subclades of H2, we also aimed to identify which SNPs are diagnostic for these subclades, when compared to the human genome reference (hs37d5). To do this we looked for segregating sites with the following properties: (1) no individual from the ingroup is ancestral at the site, (2) more than one individual from the ingroup is covered at the site, (3) no individual in the outgroup is derived at the site, and (4) more than one individual in the outgroup is covered at the site. We also restricted the search for “new” SNPs to substitutions that are not C->T or G->A, and thus are less likely to be the result of ancient DNA damage, however we included variants that are C->T or G->A in our results if they are previously-discovered SNPs in the ISOGG or YFull databases. Note, that when we report that x/N samples in a group are derived for some SNP, this means that the remaining N-x samples are not covered at this position, and we have simply recorded a “missing” base read (Supplementary Tables S5-S11).

We identified 409 potential diagnostic SNPs for the sub-haplogroups/branches in Figure 3 defined as H2 (all samples), H2d (turquoise), H2d1 (light green), H2d2 (dark green) and H2m (pastel blue), H2m1 (navy blue) and H2m2 (dark blue). Encouragingly, we found that out of the 409 diagnostic SNPs that we identified, 289 (72.6%) are already found to be H2 associated (H2-P96 or more derived) on either the ISOGG list, or on the YFull SNP list. We found only two previously discovered SNPs (0.5%), which are not associated with H2: one which is a C->T SNP associated with R1a1~ and R1a1a~ (found in 17/31 H2 samples) and a G->A SNP associated with R1b (found in 2/11 H2m2 samples). It is unlikely that the C->T substitution is due to damage since 17/31 samples have this substitution, although this is unclear for the G->A substitution. Furthermore, we also found that in our samples, which included only ancient H2 individuals (except for one modern French H2 individual), we were able to find 111 of the 134 known, basal H2 SNPs.

The remaining 110 newly discovered SNPs for the varying sub-haplogroups listed above represent either undiscovered diagnostic SNPs, or potentially lost H2 diversity (Supplementary Tables S5-S11). However, for several of our newly discovered SNPs, such as an A->G substitution at site 8611196 (found in 20/31 of our H2 samples), we find overwhelming evidence for new, true diagnostic SNPs (see Table XXX).

Our ability to detect these distinct H2 sub-haplogroups, and hence our ability to further elucidate and estimate the divergence times for an informative Y-haplogroup during the Neolithic expansion, is made possible only due to the increased coverage, and the increased number of sites we were able to target with YMCA (when compared to SG or 1240K sequencing).

## Discussion

The analysis of the Y-chromosomal history of populations can be of significant importance to the understanding of population histories. To this end we advocate for the adoption of targeted sequencing strategies for ancient Y-chromosomal sequence data. Our focused study highlighted the improved coverage and number of SNPs that are attainable when using YMCA, when compared to SG or 1240K sequencing, for the same amount of sequencing effort, accounting for endogenous human DNA content.

Targeted endogenous human DNA enrichment is of critical importance to overcome poor sample preservation in ancient DNA studies. We have shown that the Y-SNPs of the 1240K array ascertained from modern-day males, simply do not cover enough of the diagnostic SNPs on the NRY for reliable Y-haplogroup assignment, especially in the case of haplogroups that predate modern diversity highlighting a need for targeting contiguous regions in favour of an updated “Y-chromosome SNP panel”.

We were also able to show, through a deeper analysis of H2 (H-P96), that the current understanding of ancient H2 diversity is incompatible with a tree-like history (which must be true for NRY history), and that a resolution of this diversity leads to further support for the two paths of the Neolithic expansion from the Near East into Europe; an observation that would not have been possible without the improved resolution offered by YMCA.

## Materials and Methods

### Data

Note that for Y-haplogroup assignment, tree building and SNP identification, we use a mix of shotgun, 1240K, and YMCA capture sequencing runs. However, to estimate the TMRCA, we use only shotgun and YMCA data as they do not target known segregating sites (which would upwardly bias the substitution rate for samples with 1240K capture compared to those with shotgun and YMCA data only). Previously published samples were selected from published data with “H2” designated for Y haplogroups (Mathieson et al. 2015; Lazaridis et al. 2016; Lipson et al. 2017; Olalde et al. 2018, 2019; Narasimhan et al. 2019; Brunel et al. 2020; Antonio et al. 2019; Cassidy et al. 2020; Fernandes et al. 2020; Rivollat et al. 2020; Skourtanioti et al. 2020).

### Contamination quality filtering

To screen our samples for contamination, we consider the heterozygosity for sites on the NRY as our in-house samples are all merged and filtered for sites on the Y-chromosome only. We measured heterozygosity (the proportion of sites with more one than one type of base read per site) for a pileup of the quality filtered reads. We found that 45 of our 47 samples had less than 0.1%, with the remaining 2 samples 1% and 1.85% heterozygous sites. However, we were also confident in the quality of our samples as H2 is a very rare modern haplogroup, with only 19 individuals being downstream of H2-P96 on YFull at the time of this publication. Hence, if any of our samples had been contaminated by a *male* source, it would be readily noticeable in bam pileups as derived alleles for another haplogroup, which means these samples would not have been identified as H2, and hence would not be in the study.

### Method of Y Haplogroup Assignment

To assign Y haplogroups to samples we follow a partially-automated process. We begin by taking pre-prepared (i.e. trimmed, merged, deduplicated, quality-filtered) bam files, and, for each bam file, creating a pileup of every site that was covered using the *pileup* function in the Rsamtools library for the R statistical software package (Morgan 2020). We then filter this pileup of SNPs found on the ISOGG list [https://isogg.org/tree]], and then for each SNP we calculate and record the number of derived and ancestral SNP calls, the form of the ancestral and derived SNPs, and the difference (defined as the number of derived minus the number of ancestral SNP calls). Note that a positive difference indicates evidence for the ancestral form of the associated ISOGG SNP, and a negative difference indicates the converse. This method of recording the form of the called SNPs (i.e. C to T or G to A transitions), we attempt to manually identify where DNA damage infers false calls.

We return two CSVs: one CSV of only ISOGG SNPs with positive differences (for ease of reading the easiest path from root to terminal SNP), and a second CSV of all SNPs (negative or positive) allowing us to double check potentially spurious SNPs (say to check to see if more basal branches from our terminal branch are not just missing, i.e. not hit, but also not associated with negative differences). This second CSV also allows us to discover when some SNPs are derived and some are ancestral for the same branch, indicating a transitional form of the basal haplogroup.

Finally, in cases where we are uncertain of the dependability of a call (say a C to T transition with only one read), we also manually inspect where the site falls on the associated read(s), placing increased trust in SNP calls originating further from the terminal ends of a read.

### Comparing the Performance of our Y-capture Array

When comparing the empirical performance of our Y-capture array to both shotgun and 1240K sequencing, we took libraries for which shotgun, 1240K and Y-capture sequencing had all been performed. All samples were prepared and analysed using the same methods and parameters values as for the main data set.

To compare the theoretical performance of YMCA against the 1240K array, we downloaded the ISOGG SNP list v15.64. We then took the bedfiles for the NRY and 1240K array, and found which sites overlapped with the SNPs listed on the ISOGG SNP list.

When comparing empirical data performance for shotgun, 1240K and YMCA data, we included only samples that had shotgun endogenous DNA % greater than 0.1%, had sequencing results for shotgun, 1240K and YMCA sequencing and normalized the number of SNPs hit by the number of reads mapping to the human reference (hs37d5) after quality filtering. We did this to avoid any potential bias from sample quality or sequencing effort.

### Phylogenetic Tree Reconstruction

We began by taking pileups of each bam file, and performing the following quality filters for calling a consensus alignment; for each sequence we considered only sites for which we had at least two reads, with a minimum allele frequency less than 10%, and called the majority allele. We then took the aligned consensus sequences, and kept only samples for which at least 1,100 segregating sites were covered, and then filtered sites for which more than at least one sample was covered. We selected a lower bound of 1,100 segregating sites by varying this value, and inspecting bootstrap node support values. A minimum bootstrap support for major cladal splits of 80% was required.

We also included high-coverage samples from the 1000 Genomes projects (1000 Genomes Project Consortium 2015) from Y-haplogroups A, H1, H2 and H3, as well as one ancient H1 sample (Narasimhan et al. 2019) as outgroups.

We performed DNA substitution model selection using ModelFinder (Kalyaanamoorthy et al. 2017) and selected the TVM model as it had the minimum Bayesian information criterion value. We found a maximum likelihood tree using IQ-TREE v.1.5.5 (Nguyen et al. 2015).

## Data Availability

Data generated for this study can be found at the European Nucleotide Archive under the study accession number XXXXXXXXX. All R scripts can be obtained by contacting the corresponding authors. Supplementary figures and tables are available from the Dryad Digital Repository: http://xxxxxxxxx.

## Acknowledgments

We thank Nigel Bean, Jonathan Tuke, Vincent Braunack-Mayer and Gary Glonek for important discussions regarding the manuscript. We thank Chris Tyler-Smith and Yali Xue for critical advice and support during the design of the study and capture assay. This study was funded by the Max Planck Society, the French (ANR) and German (DFG) Research Foundations under the INTERACT project (ANR-17-FRAL-0010, DFG-HA-5407/4-1, 2018-2021) to M.R. and W.H., the European Research Council (ERC) under the European Union’s Horizon 2020 research and innovation program under grant agreement no. 771234 – PALEoRIDER to A.B.R., L.P. and W.H., the award Praemium Academiae of the Czech Academy of Sciences to M.E. and the project RVO 67985912 of the Institute of Archaeology of the Czech Academy of Sciences, Prague to M.S..

