## Supplementary Text for "Using Y-chromosome capture enrichment to resolve haplogroup H2 shows new evidence for a two-Path Neolithic expansion to Western Europe"

### Materials and Methods

#### Design of the YMCA Probe Set

For targeted DNA enrichment probes were designed on the basis of callable regions on the Y-chromosome as previously determined by Poznik et al. (2013). The probes were designed with a 4bp tiling and a length of 52 bp with an additional 8bp linker sequence (CACTGCGG) as described in Fu et al. (2013). Duplicated probes were removed. This resulted in 2,611,534 unique probe sequences. This probe set was spread on three Agilent one-million feature SureSelect DNA Capture Arrays. The capacity of the three arrays was filled by randomly duplicating probes from the probe set. The arrays were turned into an in-solution DNA capture library as described in Fu et al. (2013).

#### Human genome enrichment and sequencing

113 petrous bones and 52 teeth were processed in the ancient DNA laboratory at the Max Planck Institute for the Science of Human History in Jena, Germany. Upon introduction into the clean room, samples were wiped with 10% bleach and irradiated with ultraviolet light for fifteen minutes on each side. Petrous bones were either sampled by cutting them in half before drilling out bone powder from the dense portion (Pinhasi et al. 2015) or by keeping them intact and drilling into the dense portion from the outside. Teeth were sampled by removing the crown and drilling into the pulp chamber to produce bone powder. Resulting bone powder (50-100mg) was placed in 2mL Biopure tubes and stored until DNA extraction.

To the Biopure tubes containing 50-100mg bone powder one mL of extraction buffer made up of 0.9mL 0.5M EDTA, 0.025mL 0.25mg/mL Proteinase K and 0.075mL UV HPLC-water was added. The resulting mixture was incubated at 37°C for 20 hours under constant rotation. Following incubation, Biopure tubes were centrifuged at 18500 relative centrifugal force (rcf) for two minutes, separating the soluble from the insoluble parts of the mixture. The lysate was added to a 50mL falcon tube in which 10mL of binding buffer was mixed with 400µL of sodium acetate (pH 5.2, 3M). This mixture was then placed in a High Pure Extender Assembly (HPEA) tube and centrifuged at 1500 rcf for 8 minutes in a 50 mL Thermo Scientific TX-400 Swinging Bucket Rotor. Each HPEA's column was removed and inserted into a clean collection tube before centrifuging again at 18500 rcf for 2 minutes. To each column, 450µLof wash buffer from the high pure viral nucleic acid kit (HPVNAK) was added and the mixture was centrifuged for one minute at 8000 rcf. The columns were then placed into fresh collection tubes before another round of washing during which 450μL of wash buffer from the HPVNAK was added to each column followed by one minute centrifugation at 8000 rcf. Resulting columns with washed DNA were placed in 1.5 silicon tubes to which 50μL of TET was placed in the middle of the columns. The columns were incubated at room temperature for three minutes and centrifuged for one minute at 18500 rcf. Another round of adding 50μLof TET followed by incubation and centrifugation was performed and the final 100μL of DNA extract was stored at -20°C until further downstream use.

Extracts were thawed and shaken before 25μLfrom each extract was aliquoted into separate PCR strip tubes. Extracts were UDG-half treated by adding 25μL mastermix containing 0.5μL 20mg/ml BSA, 6μL 10x Buffer Tango, 6μL 10mM ATP, 3.6μL 1U USER enzyme, 0.2μL 25mM each dNTPs, and 8.7μL UV HPLC-water to each PCR strip tube. The resulting solutions were incubated for 30 minutes at 37°C and 10 minutes at 12°C. The UDG reactions were inhibited by adding 3.6μL 2U UGI to each PCR strip tube and incubating at 37°C for 30 minutes and again at 12°C for one minute. Blunt end repair was done by addition of 1.65μL 3U T4 DNA Polymerase and 3μL 10U T4 Polynucleotide Kinase, followed by incubation at 25°C for 20 minutes, and then at 12°C for 10 minutes. The resulting mixtures were purified with MinElute kits followed by elution in 20μL Elution buffer (EB) mixed with 0.05% tween. Ligation of Illumina adapters was performed through the mixture of 18μL eluate from the previous step with 1μL 5U Quick Ligase, 1μL 10μM Adapter Mix and 20μL of 2x Quick Ligase Buffer. The resulting solution was incubated for 20 minutes at 22°C and purified using a MinElute kit, followed by elution in 22μL EB containing 0.05% tween. Adapter fill in reactions were done by adding 20μL of eluate from previous step to 2μL 8U Bst 2.0 Polymerase, 0.2μL 25mM dNTPs, 4μL 10x Isothermal buffer, and 13.8μL UV HPLC-water and incubating at 37°C for 30 minutes followed by 80°C for 10 minutes. The resulting DNA libraries were stored at -20°C until indexing.

Library-specific and unique index combinations were ligated to both 5' and 3' ends of DNA fragments in each library via an indexing PCR. The total volume of each library was split into four different indexing PCR reactions which were done by mixing 2μL 10μM P5 index, 2μL 10μM P7 index, 1μL 2.5U Pfu Turbo Polymerase, 1μL 25mM each dNTPs, 1.5μL 20mg/ml BSA, 10μL 10x Pfu Turbo Buffer, 73.5μL UV HPLC-water, and 9μL of DNA library. The resulting mixture was amplified in a thermocycler with initial denaturation at 95°C for 2 minutes, followed by 10 cycles of 95°C for 30 seconds, 58°C for 30 seconds, 72°C for 1 minute, and finally 72°C for 10 minutes. The indexed libraries of the same sample were pooled and purified with a MinElute kit. Libraries were then quantified using qPCR and PCR amplified to contain 10^13 copies of DNA.

Resulting libraries were shotgun sequenced (~5,000,000 reads, either paired end with 50 cycles or single end with 75 cycles) to assess the library complexity and degree of human DNA preservation (% endogenous DNA, aDNA damage). Libraries with more than 0.1% endogenous DNA were deemed adequately preserved and selected for an in-solution hybridization enrichment (Fu et al. 2014) that targets ~10,445 kB on the NRY ("YMCA capture"). YMCA captured libraries were single end sequenced with 75 cycles to an average depth of 40 million reads per YMCA library. Libraries were not pooled prior to this capture method. After the enrichment the libraries were amplified to a concentration of 10 nM followed by a single end sequencing with 75 cycles to an average depth of 40 million reads per YMCA library.

All libraries were processed using EAGER (Peltzer et al. 2016), a modular tool that streamlines the processing of libraries from FastQC and quality filtering to mapping and duplicate removal. Sequencing adapters were clipped with AdapterRemover v2.2.0 (Schubert et al. 2016), and merged for paired-end sequencing with all reads of length <30bp discarded. The remaining reads were mapped to the human reference genome hs37d5 using BWA v0.7.1 (Li et al. 2009) with a quality filter of q30. PCR duplicates were removed using dedup v0.12.2 (Peltzer et al. 2016).

**Derivation of our method for estimating the pairwise time to most recent common ancestor (TMRCA)**

We are interested in estimating the *total* amount of evolutionary time that has passed between two samples, denoted $\tau_{ij}$, which can be separated into three non-overlapping intervals: the total evolutionary time until the last substitution occurs, and the total amount of evolutionary time that passed after the final substitution for samples *i* and *j* respectively, denoted

$$\tau_{ij} = t^*_{ij} + \delta^i_{ij} + \delta^j_{ij},$$

Respectively, where $t^*_{ij} = 2t^s_{ij} + t_{ij}$ (see Figure S1).

We begin by estimating the total evolutionary time between the i[th] and the j[th] samples, denoted $t^*_{ij}$,up until the final substitution occurred. Let $N_{ij} > 0$, be the total number of overlapping SNPs., let $K_{ij} > 2$ be the number of observed segregating sites, and let $\lambda_0 > 0$ be the rate of substitutions per site per calendar year.

We assume that each substitution occurs according to a Poisson process with rate $\Lambda_{ij} = \lambda_0 N_{ij}$, relative to the number of overlapping SNPs for individuals i and j. Hence, for some evolutionary time $t > 0$, the total number of observed substitutions has an Erlang distribution with probability density function (pdf)

$$L(\, t \mid K_{ij}, \Lambda_{ij}) \; = \frac{\Lambda_{ij} t^{(K_{ij}-1)} e^{-\Lambda_{ij} t}}{(K_{ij}-1)!}.$$

Hence, if we assume that $K_{ij}$and $\Lambda_{ij}$are known, then we may look for the optimal value of $t$ that maximizes the pdf. We do this by considering the log-likelihood function

$$l(\, t \mid K_{ij}, \Lambda_{ij}\, ) = ln\left(\frac{\Lambda_{ij}}{(K_{ij}-1)!}\right) + \left(K_{ij}-1\right) ln(t) \; - \Lambda_{ij} t.$$

Hence, we have that

$$\frac{dl(\, t \mid K_{ij}, \Lambda_{ij}\, )}{dt} = \frac{K_{ij}-1}{t} - \Lambda_{ij}.$$

If we set the first derivative of the log-likelihood to zero, we obtain a candidate maximum likelihood estimate (MLE)

$$\widehat{t_{ij}^*} = -\frac{K_{ij}-1}{t^2},$$

and since $\left(K_{ij}-1\right), t^2 > 0$, it must be that $\widehat{t_{ij}^*}$ is a maximum likelihood estimate. We also use the property that the variance of a single, unknown parameter is approximately the negative of the reciprocal of the Fisher information, e.g.

$$\hat{\sigma}^2 = \frac{K_{ij}-1}{\Lambda_{ij}^2}.$$

It must also be considered that the final substitutions probably did not occur at time $t_i$ and $t_j$, and that some time will have passed with no substitution events since the last substitution. Hence, to our MLE for the time until the final substitution $\widehat{t_{ij}^*}$, we must add some additional amount of time.

To achieve this we make the standard assumption that the time until the next substitution *would* have occurred for sample $k \in \{i, j\}$is exponentially distributed, that is, $X_{ij}^k \sim Exp(\Lambda_{ij})$. We also assume that we observed a random proportion of the interval until the next substitution, denoted $U_{ij}^k \sim U(0,1)$. Finally, we denote the amount of time that has passed in the final interval $\delta_{ij}^k = U_{ij}^k X_{ij}^k$, where $U_{ij}^k$ and $X_{ij}^k$ are statistically independent. Note then that $E[\delta_{ij}^k] = \frac{1}{2\Lambda_{ij}}$ and $Var(\delta_{ij}^k) = \frac{5/12}{\Lambda_{ij}^2}$.

Now that we have derived estimates for $t_{ij}^*$ and the $\delta_{ij}^k$, we transform these to find an estimate of the TMRCA of samples i and j, relative to the present day. Note that the true total amount of shared evolutionary time between the final substitution between samples i and j can be rewritten in terms of the shared and unshared branch times (as in Figure ??)

$$t_{ij}^* = 2t_{ij}^s + t_{ij}$$

where

$$t_{ij} = \left(t_j + \delta_{ij}^j\right) - \left(t_i + \delta_{ij}^i\right).$$

Since $E[\delta_{ij}^k] = 1/2$, it can be can be shown that

$$E[t_{ij}] = t_j - t_i \text{and } Var(t_{ij}) = \frac{1/6}{\Lambda_{ij}^2}.$$

Note that since

$$t_{ij}^* = 2t_{ij}^s + t_{ij} \Rightarrow t_{ij}^s = \frac{1}{2}\left(t_{ij}^* - t_{ij}\right)$$

a natural choice of estimator for the length of shared evolutionary time for individuals i and j would be

$$\widehat{t_{ij}^s} = \frac{1}{2}\left(\widehat{t_{ij}^*} - \widehat{t_{ij}}\right)$$

yielding an estimator for the TMRCA for individuals i and j, relative to the present day would be

$$\hat{T}_{ij} = t_i + \delta_{ij}^{i} + \hat{t}_{ij} + \widehat{t_{ij}^{s}},$$

for which a best estimator can be simplified to give

$$\hat{T}_{ij} = \frac{1}{2}\left(t_i + t_j + \frac{K_{ij}}{2\lambda_0 N_{ij}}\right).$$

Finally, we have that

$$\begin{aligned} Var(\hat{T}_{ij}) &= Var\left(t_i + \delta_{ij}^{i} + \hat{t}_{ij} + \widehat{t_{ij}^{s}}\right) \\ &= Var\left(\delta_{ij}^{i} + \tfrac{1}{2}\left[t_j + \delta_{ij}^{j} - t_i - \delta_{ij}^{i}\right] + t_{ij}^{*}\right) \\ &= Var\left(\frac{1}{2}\delta_{ij}^{1} + \frac{1}{2}\delta_{ij}^{j} + \frac{1}{2}t_{ij}^{*}\right) \\ &= \frac{1}{4}Var\left(\delta_{ij}^{1} + \delta_{ij}^{j} + t_{ij}^{*}\right) \\ &= \frac{1}{4}Var\left(\frac{5/12}{\Lambda_{ij}^{2}} + \frac{5/12}{\Lambda_{ij}^{2}} + \frac{K_{ij} - 1}{\Lambda_{ij}^{2}}\right) \\ &= \frac{K_{ij} - 1/6}{\Lambda_{ij}^{2}}. \end{aligned}$$

To test the performance of our method, we simulated 10,000 realisations of the following process. We used a grid search for the number of overlapping SNPs ranging from 20,000 to 10,000,000, (the observed values from our filtered data set) and two randomly sampled branch lengths from a log-Normal distribution from between 20,000 and 175,000 years to sample a random tree and number of overlapping sites. We used the simSeq function from the phangorn package to produce a pair of sequences (using a Jukes-Cantor model of substitution and a substitution rate of $4.5 \times 10^{-10}$substitutions per site per year per individual), from which we could count the number of pairwise segregating sites (Schliep 2011).

We found that 94.69% of the known simulated TMRCAs were within the 95% confidence intervals as calculated by our method. We found that the accuracy of our method was uncorrelated with the number of overlapping sites (p=0.951), the true TMRCA (p=0.961), or a combination of both (p=0.193) indicating that our method is unbiased for both the depth of time, and the number of overlapping sites observed in our data.

**TMRCA Estimation**

To count the number of pairwise-segregating sites between two individuals, we began by making a fasta file of aligned consensus sequences for all each bam file, and performing the following quality filter; for each sequence we considered only sites for which we had at least three reads, with a minimum allele frequency less than 10%, and called the majority allele (as suggested by Petr *et. al*. 2020) We then took the aligned consensus sequences, and calculated the number of overlapping sites for which the pair had both recorded a consensus call, and the number of pairwise segregating sites for each pair. For the substitution rate calibration we kept only pairs with >3,000,000 overlapping sites, and for the within-H2 estimates we kept only pairs with 200,000 overlapping sites, >1 segregating sites (as required by the method).

We calibrated our (relative NRY) substitution rate by fixing the mean estimated TMRCA of all Y-haplogroups A0 and all other Y haplogroups at 161,300 ybp [https://www.yfull.com]. To test if there was any effect from DNA damage or sequencing error, we first calculated a substitution rate for TMRCA estimates based on modern/modern pairs, and modern/ancient pairs. Our separate substitution estimates were within 0.687% of each other, indicating no significant increase in the substitution rate due to using ancient samples. When we estimated the substitution rate using the combined data, we found a substitution rate of $4.5 \times 10^{-10}$, which falls within the confidence intervals of existing estimates (Xue et al. 2009; Mendes et al. 2013).

**Supplementary Figures**

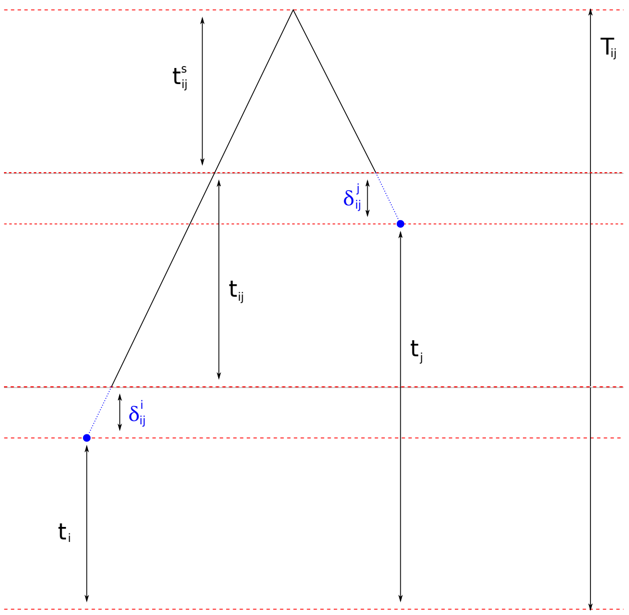


**Figure S1:** *Relative branch lengths: $T_{ij}$ is the time to most recent common ancestor ($T_{MRCA}$) for samples i and j relative to the present day, $t_i$ and $t_j$ are the calibrated ages of the samples relative to the present day, where $t_i < t_j$, $t_{ij}$ is the additional evolutionary time since the most recent common ancestor (MRCA) for sample i (compared to sample j), $t^s_{ij}$ is the shared evolutionary time per sample since the MRCA, and the $\delta^k_{ij}$ are the times between the final mutation for each lineage, and the sampling date.*

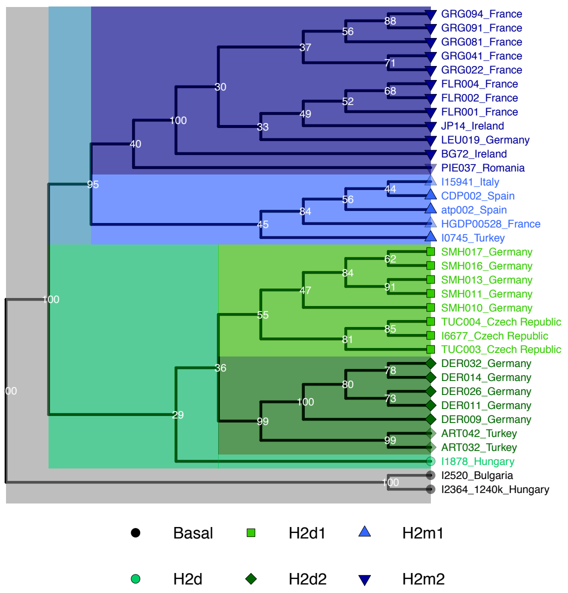


***Figure S2:*** *Phylogenetic relationships (no branch length units). Shapes and colours indicate the four major clades inferred from the phylogenetic tree. Shading indicates early to late Neolithic (solid) or post-Neolithic (transparent). Black dots indicate all non-H2 Neolithic individuals from Freeman2020 to indicate H2 sampling prevalence. Node bootstrap values are given in white.*

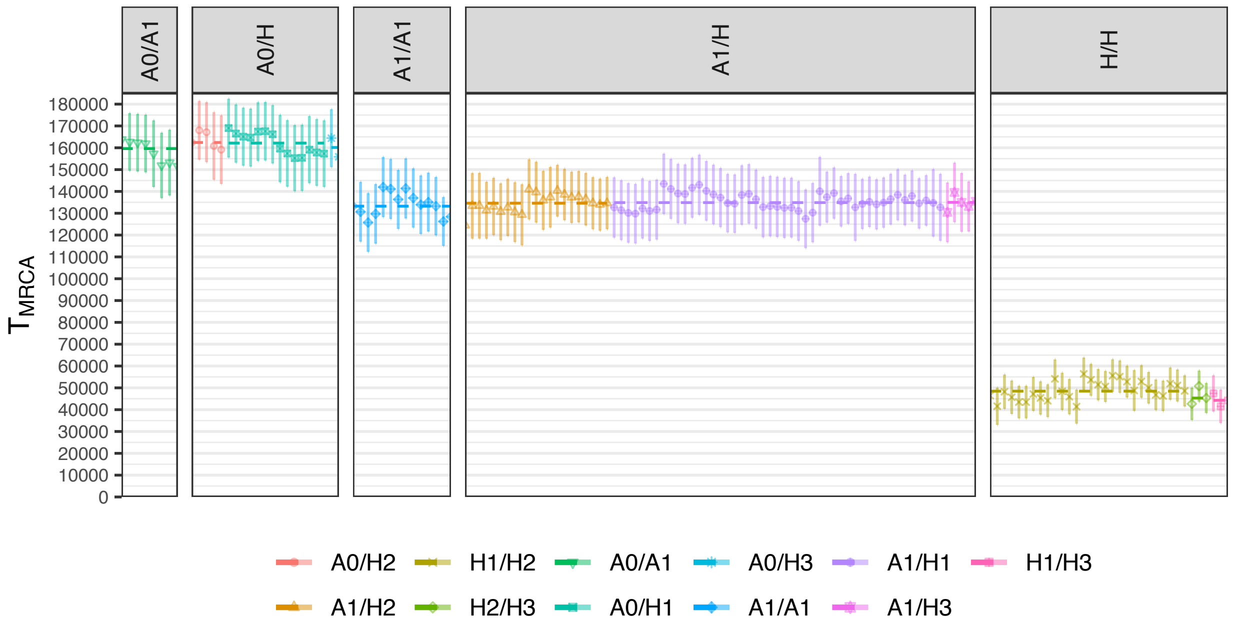


***Figure S3:*** *Estimated $T_{MRCA}$ (y-axis) for each H2 sample with pre-published individuals from Y haplogroups A0, A1, H1 and H3 (facets), calibrated by the split time of ~163 kya of A0 with all other Y haplogroups. Error bars indicated 95% confidence intervals.*

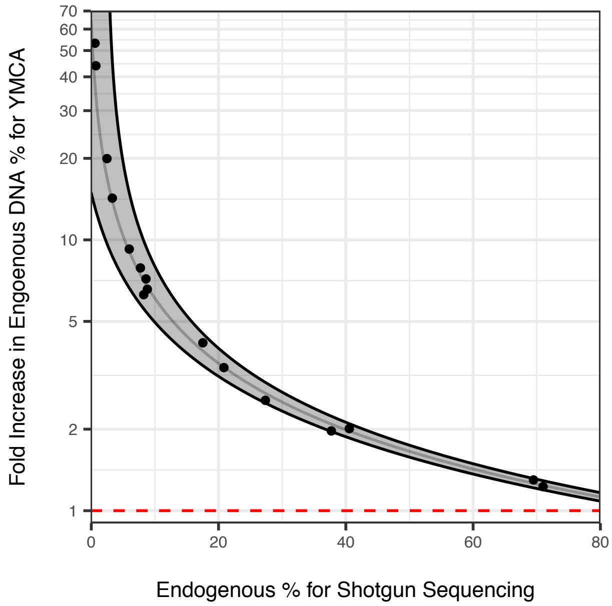


***Figure S4:*** *Fold-increase in endogenous human DNA % (y-axis) for YMCA compared to shotgun sequencing (x-axis). The shaded region indicates a 95% prediction interval, and the red dashed line indicates no improvement (a fold-increase of one).*
